# On the relation between face and object recognition in developmental prosopagnosia: Systematic association but no dissociation

**DOI:** 10.1101/034520

**Authors:** Christian Gerlach, Solja K. Klargaard, Randi Starrfelt

**Affiliations:** Department of Psychology University of Southern Denmark Denmark; Department of Psychology University of Copenhagen Denmark

**Keywords:** Developmental prosopagnosia, domain-specificity, face recognition, object recognition, selectivity

## Abstract

There is an ongoing debate about whether face recognition and object recognition constitute separate domains. Clarification of this issue can have important theoretical implications as face recognition is often used as a prime example of domain-specificity in mind and brain. An important source of input to this debate comes from studies of individuals with developmental prosopagnosia, suggesting that face recognition can be selectively impaired. We put the selectivity-hypothesis to test by assessing the performance of 10 subjects with developmental prosopagnosia on demanding tests of visual object processing involving both regular and degraded drawings. None of the individuals exhibited a dissociation between face and object recognition, and as a group they were significantly more affected by degradation of objects than control participants. Importantly, we also find positive correlations between the severity of the face recognition impairment and the degree of impaired performance with degraded objects. This suggests that the face and object deficits are systematically related rather than coincidental. We conclude that at present, there is no strong evidence in the literature on developmental prosopagnosia supporting domain-specific accounts of face recognition.

## 1. Introduction

It is currently debated whether face recognition and object recognition constitute separate cognitive domains (McKone & Robbins, 2011). Clarification of this issue can have important theoretical implications as face recognition is often used as a prime example of domain-specificity in mind and brain (Gazzaniga, Ivry, & Mangun, 2014). Domain-specificity entails the proposition that specialized cognitive functions (and brain areas) can and have evolved to handle very specific types of information; e.g., faces and faces only. As an alternative to domain-specific accounts are theories assuming the existence of more multi- or general-purpose mechanisms which can handle information from several domains; e.g., faces but also other types of objects. According to domain-specific accounts we should expect to find relatively clear-cut dissociations between domains, at least in some cases. In comparison, multi-purpose accounts would expect that information processing from different but related domains will differ in degree rather than kind.

Studies of impaired face recognition (prosopagnosia) have contributed significantly to the debate concerning selectivity of face processing. Until recently, most of these studies concerned patients with deficits in face processing following brain injury; *acquired prosopagnosia* (e.g., Damasio, Tranel, & Damasio, 1990), but now an increasing number of studies report data from individuals who have experienced face recognition problems their whole life, and where there is no (known) brain damage; a disorder known as *developmental* or *congenital prosopagnosia* (Duchaine, 2011). If face processing is the product of domain-specific operations – which are usually considered innate – we would expect that these operations can be the target of selective maldevelopment, as has been argued by for example Duchaine, Yovel, Butterworth and Nakayama (2006). If not, we would expect that object recognition is also affected in developmental prosopagnosia (DP), as has been argued by for example Behrmann, Avidan, Marotta, and Kimchi (2005).

Even though several studies of DP have been published since the seminal report by McConachie in 1976, visual object processing has typically not been assessed in great detail in these studies. Moreover, most studies have measured performance in terms of accuracy only. This may be problematic because normal performance in terms of accuracy can be achieved by means of alternative strategies that may be extraordinary time consuming and hence not normal (see e.g., Gauthier, Behrmann, & Tarr, 1999; Gerlach, Marstrand, Habekost & Gade, 2005).

Another issue concerns the way dissociations are defined. Typically, a dissociation is claimed if a person performs abnormally on task *X* but within the normal range on task *Y*, with “normality” being anchored by whether performance fall below or beyond 2 SDs of the control mean. There are two problems with this approach. First, it seems to be based on the (implicit) assumption that failure to reject the null-hypothesis regarding performance on task *Y* is proof of normality, which is a dubious assumption (Crawford, Garthwaite & Gray, 2003). Secondly, it fails to take into consideration the difference in performance between task *X* and task *Y*. In the extreme case, one could claim a dissociation if a person’s score amounted to −2.01 SD on task *X* and −1.99 SD on task *Y*; a trivial difference of .02 SD. These problems are not specific to the studies of DP but have been discussed in other contexts as well (see e.g., Laws, 2005). To avoid them Crawford et al. (2003) have suggested that a performance pattern must fulfil two criteria in order to count as a dissociation: (i) the person’s performance on task *X* must differ significantly from that of the normal population, and (ii) the *difference* in performance of that person on task *X* and *Y* must differ significantly from the difference-scores of the normal population on task *X* and *Y*. To our knowledge such a “qualified” dissociation has not yet been reported in DP, or for patients with acquired prosopagnosia for that matter.

In summary; while there have been many reports of dissociations between face and object recognition performance in individuals with DP, no study has yet demonstrated such a dissociation in terms of both accuracy and reaction time while also adopting the stringent criteria for a dissociation advocated by Crawford et al. (2003).

In the present study we tested 10 individuals with DP to examine if any of them would fulfil the criteria for a dissociation as described above, that is: (i) performing worse with faces than with objects in terms of both accuracy and reaction time, and (ii) satisfying the criteria for a classical or strong dissociation as defined by Crawford et al. (2003). We examined this by means of three different types of visual object processing tasks. The first type of task was *object decision*, where participants must decide whether a displayed object is real or nonsense. All non-objects were chimeric combinations of real objects (for example, front part horse and back part dog). The use of chimeric non-objects makes the task more demanding in terms of perceptual differentiation than for example object naming tasks (Gerlach et al., 2005; Gerlach & Toft, 2011). To further increase the sensitivity of the task, we presented the stimuli as silhouettes and fragmented forms in addition to presenting them as regular line drawings. The use of degraded stimuli (e.g., fragmented forms) has previously proven successful in revealing even subtle deficits in visual object processing which may otherwise go unnoticed (Starrfelt, Habekost & Gerlach, 2010). The second type of task involved within-class recognition of objects (cars). In this task, the participants must be able to keep a stimulus in memory for a short duration and compare it with an array of objects that, just like faces, are very similar to each other and the target. The third type of task required perceptual matching of two simultaneously presented stimuli; either two faces or two houses. Hence, as opposed to the two other types of task, this task does not rely on memory to any great extent. Stimulus-pairs could differ in either 1^st^ order relations among the elements (e.g., the nose placed below the mouth in one face but at its normal position in the other), 2^nd^ order relations among elements (e.g., difference in the spacing between the eyes), or in their features (e.g., different shape of the nose); for a discussion of these dimensions in face processing see Maurer, Le Grand, & Mondloch (2002). Furthermore, the number of differences between the stimulus-pairs varied parametrically from one to four differences along the three dimensions. Hence, with this design we could examine if the DPs were impaired in perceiving particular sorts of information (1^st^ order, 2^nd^ order and/or features), if their problems were modulated by visual similarity (degree of difference), and if potential difficulties were specific to faces.

If face recognition can be selectively impaired in DP, we expect our sample, or at least some of the individuals with DP, to perform within the range of control subjects on these tasks, and further, that the discrepancy in their performance with faces compared to objects be significantly larger than that observed in control subjects (cf. Crawford et al., 2003).

## 2. Assessment of face recognition ability

### 2.1. Method & Subjects

All participants provided written informed consent according to the Helsinki declaration to participate in the project. The Regional Committee for Health Research Ethics in Southern Denmark has assessed the project, and found that it did not need formal registration.

Following some appearances in Danish media, where we have informed about developmental prosopagnosia, we have been contacted by a number of people complaining of face recognition problems. They all report difficulties recognizing friends, colleagues, and sometimes even close family members and themselves by their faces, and that these problems have been present throughout their life.

10 of these subjects are included in the current study, along with 20 controls matched for age, gender and education. As there are no established diagnostic criteria for DP we initially included subjects in our sample if they performed abnormally on the Cambridge Face Memory Test (CFMT; Duchaine & Nakayama, 2006) and/or the Cambridge Face Perception Test (CFPT; Duchaine, Germine, & Nakayama, 2007) when compared to the age and gender adjusted norms provided by Bowles et al. (2009). These tests were kindly provided by Brad Duchaine and translated into Danish. In the current study, only participants who performed more than 2SD’s below the mean of the matched control group on the CFMT, the most commonly used diagnostic test, were included. All included DP’s also report severe difficulties with face recognition in their everyday life, as evaluated by a 29-item systematic questionnaire using a 4-point Likert scale (Freeman, Palermo, & Brock, 2015). This questionnaire was kindly provided by Romina Palermo and Jon Brocks, and translated to Danish by the first author. On the CFPT, six of the DPs scored significantly below the mean of the control group. In comparison, all DPs performed within the normal range on The Autism-Spectrum Quotient (AQ) questionnaire (Baron-Cohen, Wheelwright, Skinner, Martin, & Clubley, 2001). See Table 1 for an overview of age, gender, and basic test scores. The DP subjects did not receive remuneration for their participation in this study.

**Table 1.** Age, gender and performance on the Cambridge Face Memory Test (CFMT), the Cambridge Face Perception Test (CFPT), the Face Questionnaire (FQ), and the Cambridge Car Memory Test (CCMT) for the 10 individuals with developmental prosopagnosia. The mean performance and SD for the controls’ scores are also listed. Values in boldface designate performance deviating significantly from the mean of the matched control group as assessed by means of Crawford, Garthwaite & Porter’s (2010) method. In the CFMT a low score indicates a deficit, while in the CFPT and in the FQ a high score indicates a deficit.

| Case | Age | Gender | CFMT | CFPT | FQ | CCMT |
| --- | --- | --- | --- | --- | --- | --- |
| PP04 | 57 | Male | **37** | **86** | **71** | 42 |
| PP07 | 40 | Female | **41** | 60 | **66** | 55 |
| PP09 | 40 | Female | **43** | **70** | **52** | 47 |
| PP10 | 34 | Female | **33** | 58 | **62** | 45 |
| PP13 | 51 | Male | **35** | 42 | **64** | 39 |
| PP16 | 23 | Female | **39** | **64** | **54** | --- |
| PP17 | 49 | Female | **35** | **88** | **56** | **33** |
| PP18 | 38 | Female | **30** | **78** | **69** | 41 |
| PP19 | 16 | Male | **33** | 48 | **53** | 40 |
| PP27 | 25 | Male | **42** | **66** | **59** | 45 |
| Control mean |  |  | 59.1 | 41.3 | 22.4 | 50.9 |
| Control SD |  |  | 7.9 | 11.4 | 11.4 | 7 |

## 3. Assessment of visual object recognition ability

DPs and controls were tested on three object decision tasks which varied in stimulus type (regular line drawings, silhouettes, and fragmented forms). The demand placed on perceptual differentiation in object decision can be controlled by manipulating the type of nonobjects used. If the nonobjects are novel, that is, completely unknown to the participants, the task is relatively easy, as the only judgement needed is one of familiarity. In the present case the nonobjects were partly familiar because they were chimeric nonobjects composed by exchanging single parts belonging to different real objects. As demonstrated in previous studies, it is harder to reject chimeric nonobjects as being real objects than to reject novel nonobjects as being real objects (Gerlach, Law, Gade, & Paulson, 1999; Gerlach, 2001). This effect of task difficulty also affects the processing of the real objects presented: It is harder to recognize a real object as a real object when it is presented in the context of chimeric nonobjects than when it is presented in the context of novel nonobjects. This can be explained in the following way: Given that the nonobjects in the easy tasks are novel it might be sufficient to identify just a few recognizable parts of an object to judge it as a real object. This strategy will not suffice in the difficult task because the nonobjects are composed of parts of real objects. Accordingly, when discriminating between real objects and chimeric nonobjects, it is necessary to keep on processing until a particular representation in visual long term-memory wins the competition and a complete match is found, i.e. the object is recognized. Otherwise, one will risk judging a nonobject as a real object (Gerlach & Toft, 2011).

### 3.1. Method

#### 3.1.1. Design

The DPs and the 20 control participants performed all three object decision tasks in the same order (fragmented forms, full line drawings, and silhouettes), except for PP16 who did not perform the version with silhouettes.

In each task, the participants were instructed to press “1”, on a serial response box, if the picture represented a real object and “2”, if it represented a nonobject. Participants were encouraged to respond as fast and as accurately as possible. Prior to each of the three tasks, the participants performed a practice version of the upcoming task. Stimuli used in these practice versions were not used in the actual experimental conditions.

The DPs and the control participants performed the fragmented version and the full line drawing version on the same day. The silhouette version was performed on another occasion separated by at least two weeks (mean = 211 days, range 20 – 274 days) for the DPs and one week for the control participants (mean = 109 days, range 8 – 199 days). Although this difference in interval was significant (*t* (1, 26) = 3.91, *p* < .001), there was no evidence that a shorter interval had a positive impact on performance with the silhouette version. Hence we failed to find any significant negative correlation between interval and discrimination sensitivity *(A)*, or a significant positive correlation between interval and RT for neither the DPs nor the Controls (all *p*’s > .4 one-tailed). Accordingly, in the case that the group of DPs perform worse than the control group with the silhouette version, this performance difference cannot be attributed to a difference in test interval.

#### 3.1.2. Stimuli

160 pictures were presented in each task: 80 real objects and 80 chimeric nonobjects. The full line drawings of real objects were taken from the set of Snodgrass and Vanderwart (1980). The 80 chimeric drawings of nonobjects were selected mainly from the set made by Lloyd-Jones and Humphreys (1997). These nonobjects are line-drawings of closed figures constructed by exchanging single parts belonging to objects from the same category (see figure 1).

The fragmented versions of the regular line drawings were made by imposing a mask as a semi-transparent layer on the regular line drawings. This mask consisted of blobs of different sizes and shapes. The regular line drawing and the mask were subsequently merged into a single layer yielding a fragmented version of the regular line drawing (see figure 1). The same mask was used for the generation of all fragmented stimuli. The silhouette versions of the regular line drawings were made by replacing the colour of each pixel within the interiors of the regular line drawings with the colour black (see figure 1). The order of pictures was randomized within each task.

**Figure 1.**
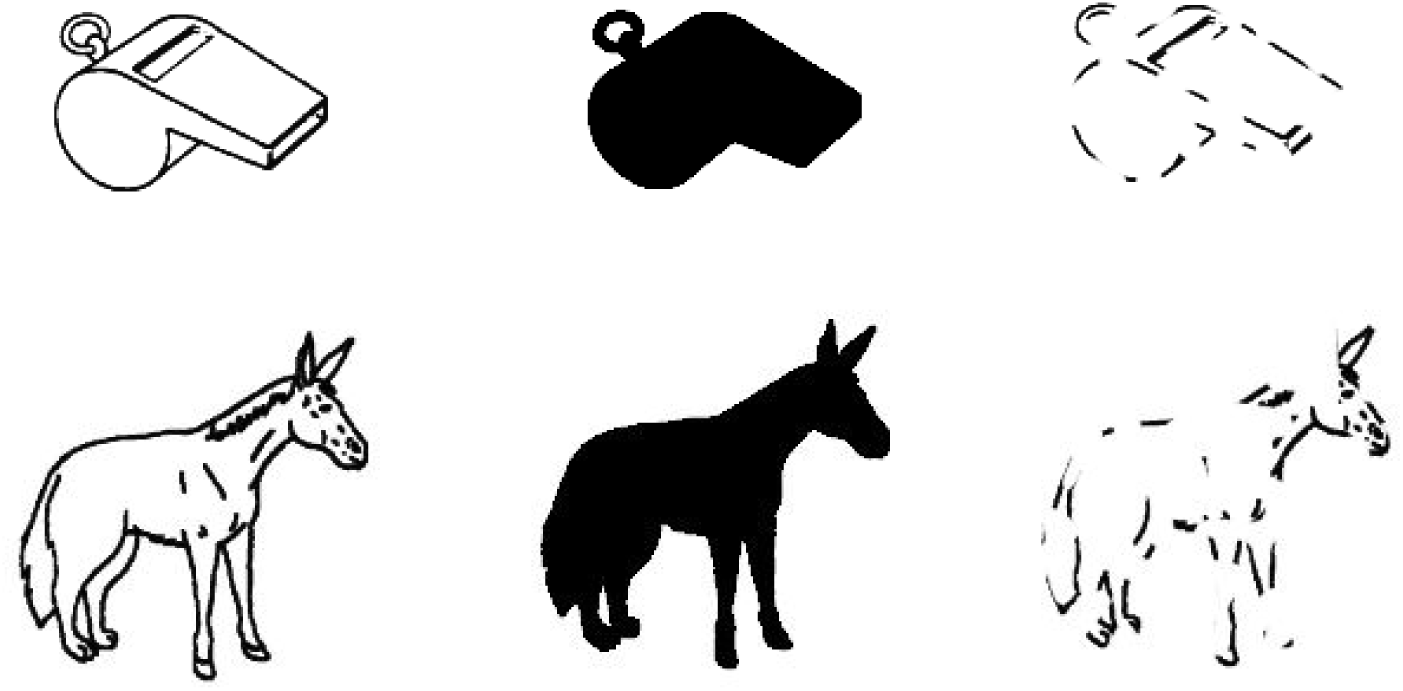
Examples of the stimuli used in the Object Decision Tasks. Upper panel: three versions (full drawing, silhouette, and fragmented) of a real object. Lower panel: three versions of a chimeric nonobject.

#### 3.1.3. Procedure

All stimuli were presented centrally in black on a white background on a PC-monitor and subtended 3–5° of visual angle. The stimuli were displayed until the participant made a response. The interval between response and presentation of the next object was 1 s. RTs were recorded by use of a serial response box.

#### 3.1.4. Statistical analyses

We used *A* as a measure of discriminability (Zhang & Mueller, 2005). This is a bias-free measure that varies between 0.5 and 1.0 with higher scores indicating better discrimination between objects and nonobjects. Prior to analyses of *A*-scores we screened for individuals exhibiting extreme hit/false alarm rates (rates approaching 1), as such cases may yield unreliable estimates of discriminability. One subject (PP04) had such a performance on the object decision task with silhouettes where his hit rate was .99 while his false alarm rate was .94 (indicating that he pressed 1 for almost all trials). Even though this subject clearly had difficulties with this task – indeed his performance was on chance level (53% correct responses) – we decided to exclude him, and his matched controls, from the analyses of the results from this task as his *A*-score and RTs could not be clearly interpreted. No other subjects were excluded on this account.

All analyses of RTs presented below are based on correct responses to real objects only, as the nonobjects served no other purpose in the present study than to ensure detailed shape processing of the real objects. Prior to RT-analysis the data were trimmed excluding trials from a particular participant if the RT of that trial fell above or below 2.5 SD of the mean of the participant’s RT. For no subject in any of the three tasks did trimming result in discard of more than 5% of the trials. Comparisons between the DPs and the control participants were based on non-parametric tests as the variables departed from normality.

### 3.2. Results

The comparisons between the scores of the DPs and the control group were based on Mann–Whitney *U* exact test. As can be seen in Table 2, the DPs performed within the normal range with regular drawings. In comparison, the DPs performed significantly worse than the control group with silhouettes in terms of *A* and marginally so in terms of RT. In the condition with fragmented drawings they were marginally slower than the controls but did not differ significantly from the controls in terms of *A*.

**Table 2.** Statistics (Median, interquartile range (IRQ,), Z-score, and p-value) associated with the comparisons of the DP group and the control group in the three object decision tasks. The comparisons are based on Mann–Whitney U exact test.

| *Condition* | DPs |  | Controls |  | *Z* | *p*-value |
| --- | --- | --- | --- | --- | --- | --- |
|  | *M* | *IRQ* | *M* | *IRQ* |  |  |
| Regular drawings *A*: | .945 | .025 | .958 | .026 | −0.84 | .42 |
| Regular drawings RT (ms): | 860 | 263 | 707 | 226 | −1.23 | .23 |
| Silhouette drawings *A*: | .873 | .068 | .926 | .05 | −2.33 | .02 |
| Silhouettes drawings RT: | 1189 | 590 | 899 | 277 | −1.77 | .08 |
| Fragmented drawings *A*: | 784 | .086 | .800 | .073 | −0.6 | .56 |
| Fragmented drawings RT: | 1135 | 431 | 922 | 384 | −1.94 | .06 |

### 3.3. Single subject analysis

To examine the performance of the DPs individually, we compared the scores *(A* and mean RT) of each DP with the mean score of the control participants using the method developed by Crawford, Garthwaite and Porter (2010) for comparison of an individual’s score with that of a small control sample. This was done for all three object conditions. As can be seen in Table 3, where the results of these comparisons are summarized, three of the DPs (PP07, PP16 & PP27) scored within the normal range in terms of both *A* and RT in all conditions. Especially PP07 and PP27 seemed to perform quite well. In section 6.0 we examine the question of whether any of these normally performing DPs exhibit a classical/strong dissociation in accordance with criteria suggested by Crawford et al. (2003).

**Table 3.** A -scores and RT (ms.) for each of the participants with developmental prosopagnosia on the three object decision tasks. Values in boldface designate performance deviating significantly from the mean of the matched control group as assessed by means of Crawford, Garthwaite and Porter’s (2010) method.

| Case | Regular drawings |  | Silhouettes |  | Fragmented |  |
| --- | --- | --- | --- | --- | --- | --- |
|  | *A* | RT | *A* | RT | *A* | RT |
| PP04 | .965 | 1025 | ----- | ----- | .782 | 1303 |
| PP07 | .941 | 584 | .894 | 638 | .786 | 730 |
| PP09 | .973 | 706 | .905 | **1251** | .882 | 947 |
| PP10 | .945 | 895 | **.833** | **1450** | .703 | 1426 |
| PP13 | .955 | 895 | .868 | **1260** | .773 | 953 |
| PP16 | .963 | 719 | ----- | ----- | .853 | 1368 |
| PP17 | **.865** | 672 | .903 | 728 | .767 | 1014 |
| PP18 | .945 | **2044** | **.805** | **2230** | **.694** | **2475** |
| PP19 | .936 | 939 | **.847** | 1127 | .820 | 1234 |
| PP27 | .940 | 825 | .930 | 1065 | .831 | 1037 |
| Control mean | .951 | 794 | .914 | 899 | .795 | 1002 |
| Control SD | .024 | 230 | .03 | 184 | .057 | 292 |

### 3.4. Within-class object recognition

To examine within-class object recognition performance, nine of the 10 subjects with DP were tested on the Cambridge Car Memory Test (Dennett et al., 2011) (PP16 did not take this test, and hence her matched control participants were also not included in the analysis), which is a test equivalent to the Cambridge Face Memory Test using cars as stimuli. The maximum score in this test is 72.

#### 3.4.1 Results

The median score of the DPs was 42 (IQR = 6.5). In comparison the median score of the control subjects was 51 (IQR = 10.3). This difference was significant (Mann–Whitney *U* exact test, Z = −2.6, *p* = .008). Analysis at the single subject level by means of Crawford, Garthwaite and Porter’ method (2010) revealed that only PP17 scored outside the range of the control subjects (see table 1).

### 3.5. Discussion

The results presented above suggest that our sample of DPs performed within the normal range on a quite demanding task of visual object recognition with regular line drawings. When the same task involved degraded stimuli, subtle deficits were nevertheless revealed. Hence, with silhouette drawings the group of DPs were significantly impaired relative to the control participants in terms of discriminability and also performed more slowly although this difference in latency was only marginally significant. With fragmented forms the DPs also exhibited prolonged RTs; again an effect that was only marginally significant. When we consider these deficits subtle, it is because three out of eight of the DPs performed within the normal range with silhouettes in terms of both discriminability and RT, whereas nine out of 10 performed within the control range with fragmented forms. Indeed two of the DPs fell within the performance range of the control sample in all three tasks in terms of both RT and discriminability. A similar pattern was seen for performance on the CCMT. Here the DP group scored significantly below the control group even though only one DP fell outside the range of the control subjects based on single case statistics.

## 4. Relating face and object recognition abilities

So far we have established that the group of DPs performed abnormally with object recognition of degraded stimuli and within-class recognition of objects (cars). This, however, does not indicate that these deficits are directly related to the face processing deficits observed; it could reflect associated deficits. Moreover, one may even question the validity of using impoverished stimuli in the first place.

In many instances where the demand on perceptual differentiation is low, successful classification of an object can be based on a limited amount of information, e.g., a couple of features (Gerlach et al., 2005; Gerlach & Toft, 2011). This is the reason why we chose to use a difficult object decision tasks. The decision to also use impoverished stimuli is a logical extension of this line of reasoning because subtle deficits may only become apparent when the recognition system is challenged (Starrfelt, Habekost & Gerlach, 2010). We note that similar assumptions lie behind the construction of the Cambridge Tests in that they also make use of impoverished stimuli (faces and cars with added noise). It is also the case that the object recognition system often deals with stimuli that are impoverished for natural reasons. Objects may be partly occluded by other objects (causing them to appear fragmented), or they may be viewed in dim light or strong backlight (causing them to appear as silhouettes). Hence, the use of fragmented forms and silhouettes per se does not necessarily make the recognition situation artificial (or un-ecological).

While the use of impoverished stimuli can be justified, and even commendable, the observed association between face and object recognition performance can still be considered coincidental. To show that the connection is more than coincidental, a systematic relationship between variation in object recognition performance and variation in face recognition performance must be established. We examine this possibility below.

### 4.1. Procedure

To examine whether variation in object recognition performance is systematically related to face recognition performance, we compared the scores obtained from the CFMT with scores from the CCMT and the discriminability scores obtained from the two object decision conditions where there were signs of impairment; the one with silhouettes and the one with fragmented forms. This was done by means of correlation analyses. We did not perform the same analyses with the RTs from the object decision tasks with degraded stimuli because the CFMT is based on accuracy only. We report statistics derived by means of Pearson's correlation coefficient (*r*). These analyses were performed separately for the DPs and control participants.

### 4.2. Results

There was a significant positive correlation between performance on the CFMT and *A*-scores with silhouettes (*r* = .87, *p* = .005, 95% CI = [.55, 1] based on bootstrap analysis with1000 samples) for the DP group but not for the control group (*r* = -.18, *p* = .51). There was also a significant positive correlation between performance on the CFMT and *A*-scores with fragmented forms (*r* = .78, *p* = .007, 95% CI = [.26, .98] based on bootstrap analysis with1000 samples) for the DP group but again not for the control group (*r* = -.04, *p* = .87). The correlation between the CFMT and the CCMT failed to reach significance in both the DP (*r* = .58, *p* = .1) and the control group (*r* = .07, *p* = .76). For a graphical illustration of the significant findings see Figure 2.

**Figure 2.**
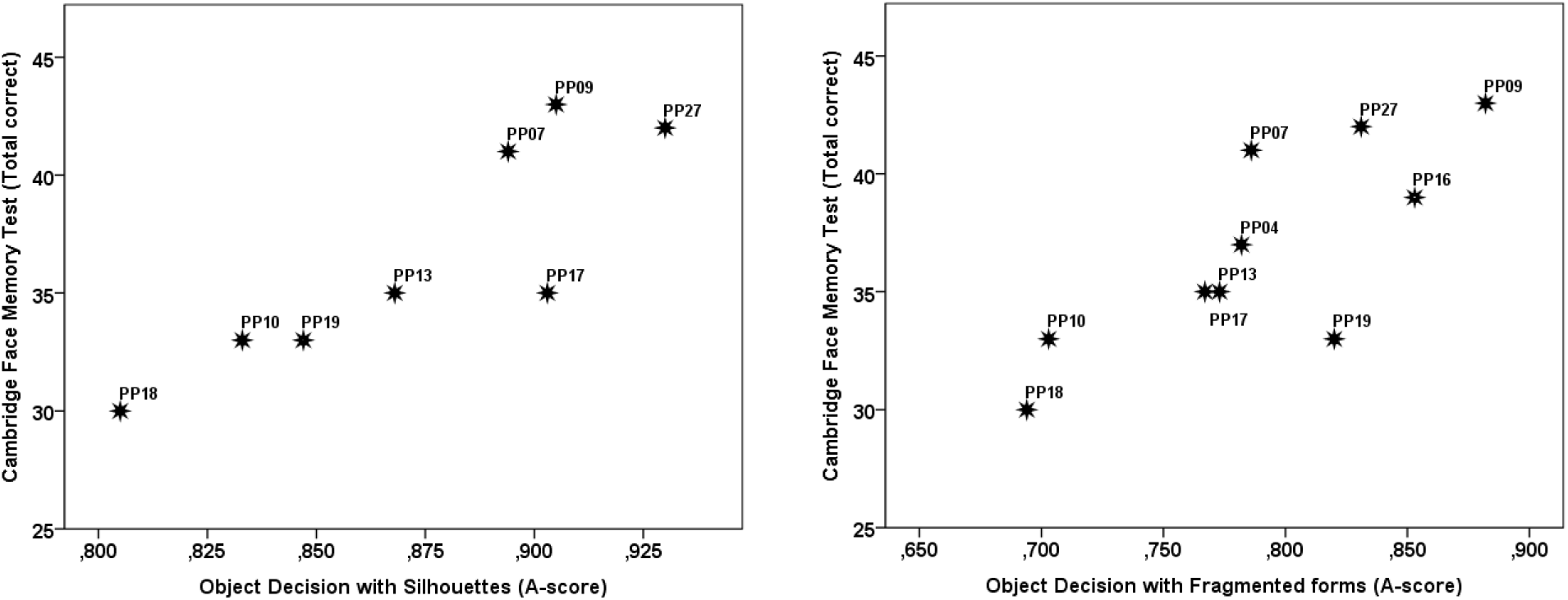
Scatterplots showing the relationship between performance on the Cambridge Face Memory Task and the object decision tasks with silhouettes (left panel) and fragmented forms (right panel) in the DPs.

### 4.3. Discussion

For the DP-group, performance on the CFMT correlated significantly with performance on both the object decision task with silhouettes and the object decision task with fragmented forms. While caution should be exercised in interpreting correlations based on such small samples *(n* = 8 for silhouettes and *N* = 10 for fragmented forms), we note that the correlations were quite reliable as reflected by the fact that the 95 % CIs did not include 0. Accordingly, while there is some uncertainty associated with using the correlations as estimates of correlations in the general population of DPs, they do seem impressive in that the lower bound was *r* = .55 for object decision with silhouettes and *r* = .26 for fragmented forms. These findings suggest that the face and object recognition deficits observed in the DP group are systematically related and thus unlikely to reflect two associated deficits.

## 5. Perceptual matching

### 5.1.1. Subjects

All DPs except for PP16 participated *(n* = 9). To keep the DP group and the Control group matched we excluded the two controls of PP16 yielding a total of 18 subjects in the control group.

### 5.1.2. Design

The subjects were presented with two stimuli at a time, either two faces or two houses, and had to decide whether the stimuli were identical or differed. They were instructed to press the 'same'-key on a serial response-box (index finger) if the stimuli were identical and the 'different'-key (middle finger) if the stimuli differed in any respect. The stimuli could differ in either 1^st^ order relations (e.g., the nose placed below the mouth in one face but at its normal position in the other), 2^nd^ order relations (e.g., difference in the spacing between the eyes), or in their consistent features (e.g., different shape of the nose). Furthermore, the differences along these three dimensions varied parametrically between the stimulus-pairs as illustrated in Figure 3.

**Figure 3.**
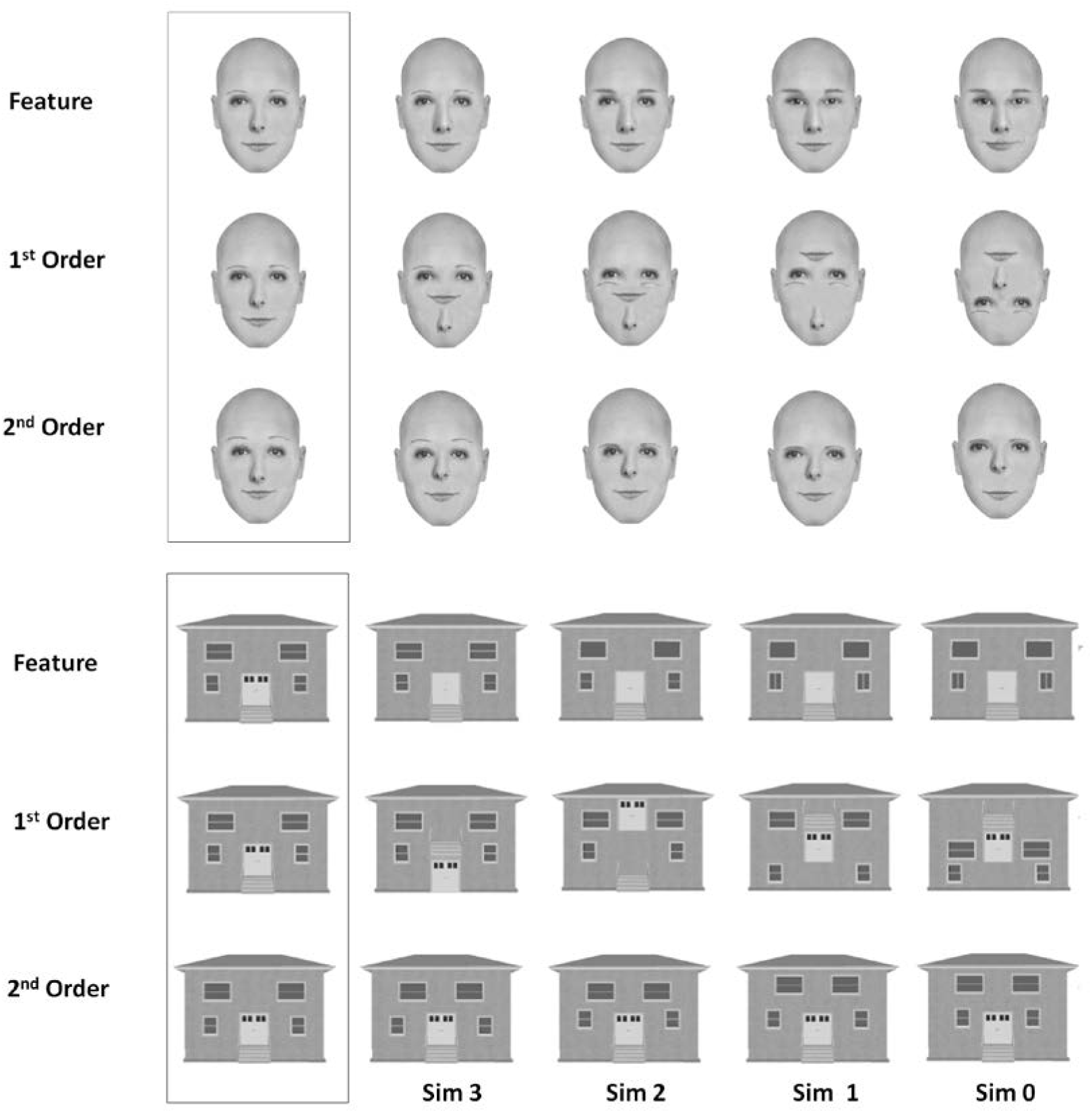
Examples of the face and house stimuli used in the perceptual matching task. “Sim” designates similarity level with each stimulus differing from the one presented in the boxed area by 1 difference (Sim 3), 2 differences (Sim 2), 3 differences (Sim 1), and four differences (Sim 0).

In the task instructions the subjects were made aware that there was an equal amount of identical and different stimulus-pairs. They were also informed that the stimuli could differ in either 1^st^ order relations, 2^nd^ order relations or in their constituent features. These differences were also illustrated by example stimulus-pairs which were not used in the actual experiment. Finally they were told that the stimulus-pairs could differ in how similar they were; as an example, some pairs would differ in only one feature whereas others would differ in several of their constituent features. The subjects were encouraged to respond as fast and as accurately as possible. Prior to the actual experiment the subjects performed 48 practice trials with all combinations of differences (1^st^ order, 2^nd^ order and featural) and similarity levels. Stimuli used in practice trials were not used in the actual experimental conditions. Feedback was provided during practice but not during the actual experiment. A similar experimental paradigm, using the same stimuli as the present, has previously been used by Collins, Zhu, Bhatt, Clark and Joseph (2012) and by Joseph, DiBartolo and Bhatt (2015). The stimuli used in the present experiment were kindly made available by Jane E. Joseph. Hence, for specification of stimulus parameters, beyond what is given below, we refer to these studies.

### 5.1.3. Stimuli

A total of 560 stimulus-pairs were presented. 176 were 1^st^ order stimulus-pairs [11 different face-pairs by 4 different similarity levels + 11 different House-pairs by 4 different similarity levels + 11 identical face-pairs by 4 different combinations + 11 identical house-pairs by 4 different combinations], 192 were 2^nd^ order stimulus-pairs [12 different face-pairs by 4 different similarity levels + 12 different House-pairs by 4 different similarity levels + 12 identical face-pairs by 4 different combinations + 12 identical house-pairs by 4 different combinations], and 192 were featural stimulus-pairs [12 different face-pairs by 4 different similarity levels + 12 different House-pairs by 4 different similarity levels + 12 identical face-pairs by 4 different combinations + 12 identical house-pairs by 4 different combinations]. The stimulus-pairs were arranged such that the two faces/houses were presented one above the other. Each stimulus in a pair subtended 2.5–5.3^°^ of visual angle.

### 5.1.4. Procedure

Each stimulus-pair was shown until the subject made a response or for a maximum duration of 10s. If no response was made within 10s the trail was terminated and counted as an error. The interval between response (or termination) and presentation of the next stimulus-pair was 1.5s. The presentation order of the 560 stimulus-pairs was randomised with a rest period inserted following a row of 56 trials. The rest period was terminated by the subject when s(he) was ready to proceed.

### 5.1.5. Statistical analyses

We measured overall discriminability between same and different trials in terms of *A*. Prior to analyses of *A*-scores we screened for individuals exhibiting extreme hit/false alarm rates (rates approaching 1). No such cases were observed. While *A* can be computed for faces and houses separately over all conditions, they cannot also be computed separately for the 1^st^ order, 2^nd^ order or featural condition because stimulus presentation was randomised such that stimulus-pairs with identical faces/houses (same-responses) could not be assigned to a particular condition (1^st^ order, 2^nd^ order or featural). Furthermore, as accuracy was quite low, and for some subjects at chance-level in some of the conditions (see below), we decided not to perform a full analysis of the RT-data. Finally, given that we were interested in effects of visual similarity, a dimension that only varied for different-trials, analyses of accuracy data were limited to different-trials only (number of correct responses). Comparisons between the DPs and the control participants were based on non-parametric tests as several of the variables of interest departed from normality.

### 5.2. Results

#### 5.2.1. Discriminability

The median *A*-score of the DPs for faces was .932 (IQR = .192) and .978 (IQR = .029) for the control group. This difference was significant (Mann–Whitney U exact test, Z = −2.5, *p* = .011). The median *A*-score of the DPs for houses was .939 (IQR = .1) and .965 (IQR = .02) for the control group. This difference was also significant (Mann–Whitney U exact test, Z = −2.8, *p* = .004).

#### 5.2.2. Accuracy

To examine whether the DP group generally performed differently than the control group on 1^st^ order relations, 2^nd^ order relations and featural differences for faces and houses, we first computed the mean number of correct responses to different-trials averaged over the four similarity levels for each subject for each of the six conditions. We next compared these median scores of the DP group and the control group for the six conditions by means of Mann–Whitney *U* Exact test. As can be seen from table 4, where these results are summarized, the DP group differed from the control group in all conditions except for 1^st^ order differences in faces. Although the range also differed between the groups, the variability between them did not differ significantly (Moses Test of Extreme Reaction; all *p*’s > .05). This suggests that the differences observed reflect differences in the medians rather than in the distributions of scores.

**Table 4.** Comparison of the median number of correct responses to different-trials (range in brackets), averaged over the four similarity levels, for the DP group and the control group. Differences between groups were examined by means of Mann–Whitney U Exact tests.

| Condition | DP group | Control group | Z | *p*-value |
| --- | --- | --- | --- | --- |
| 1st order faces | 44 (13-44) | 44 (33-44) | -.84 | .41 |
| 1st order houses | 43 (1-44) | 44 (41-43) | -2.0 | .05 |
| 2nd order faces | 32 (1-44) | 45 (16-48) | -2.96 | .002 |
| 2nd order houses | 26 (0-44) | 39 (13-47) | -2.63 | .007 |
| Featural faces | 39 (10-46) | 46 (37-48) | -2.56 | .009 |
| Featural houses | 38 (11-43) | 42 (31-47) | -2.82 | .004 |

#### 5.2.3. Effects of visual similarity

To examine effects of visual similarity in the six conditions we first computed the mean percentage of correct responses for each similarity level across the subjects within each group. We then computed the correlation between similarity level and the mean percentage of correct responses to different-trials for each of the six conditions for each of the two groups by means of Pearson's correlation coefficient. Note that these analyses ignore intersubject variability because they are based on the grand mean of all subjects across a particular similarity level and not on the means of the individual subjects for a particular similarity level. Accordingly, the degrees of freedom for each analysis were 4. The correlation analyses were based on one-tailed statistics as we had a directional hypothesis: Accuracy will decrease as similarity increase. As can be seen from figure 4, where the results of these analyses are summarised, accuracy generally did decrease as similarity increased. However, in 5 out of 12 cases this linear effect was not significant: 1^st^ order differences in faces (Controls); 1^st^ order differences in houses (DPs and Controls), 2^nd^ order differences in houses (DPs and Controls).

**Figure 4:**
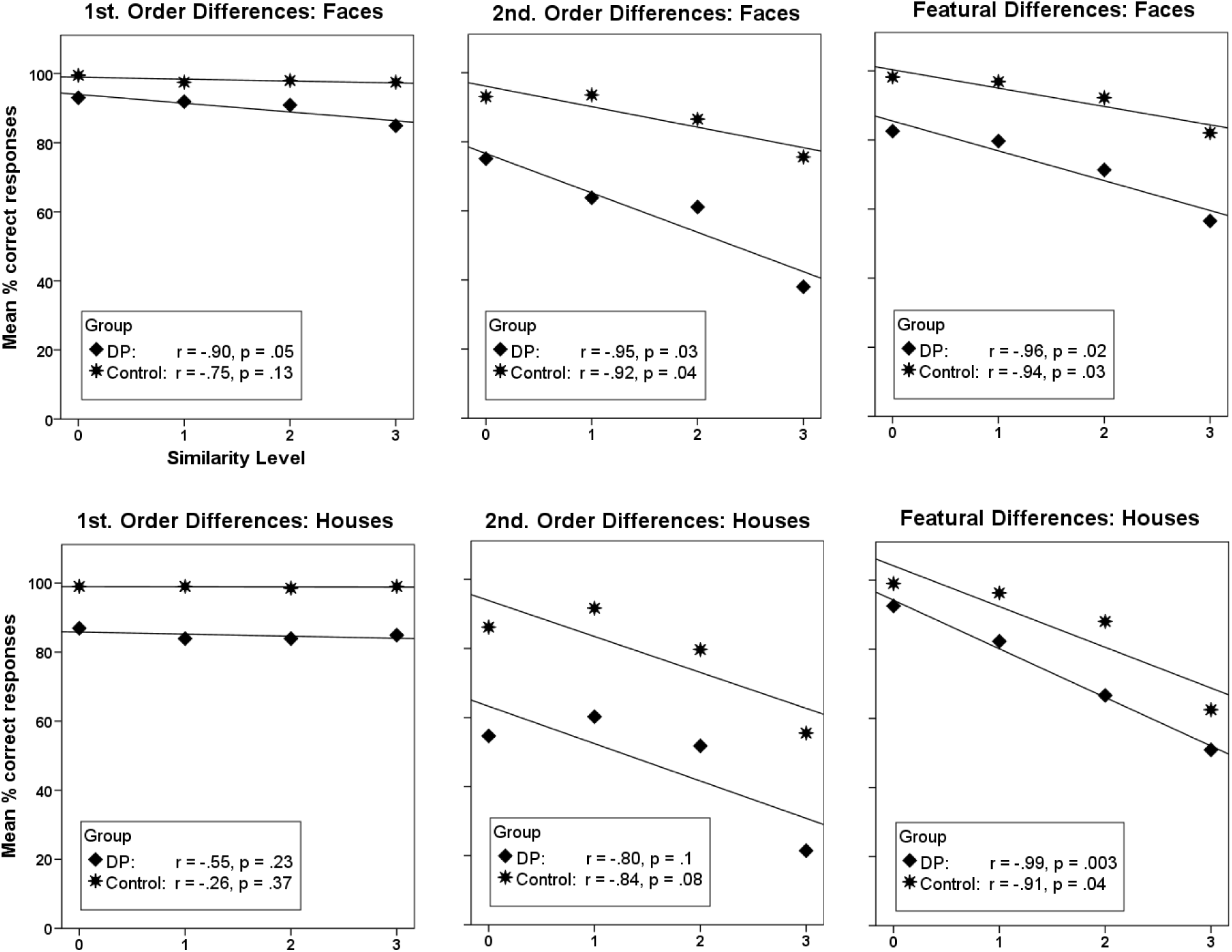
Scatterplots showing the relationship between similarity level and mean % correct responses to different trials in the six conditions of the perceptual matching experiment for the developmental prosopagnosics (DP) and the control group. Similarity increases parametrically from level 0 to level 3, with 0 indicating minimum similarity with four differences and 3 indicating maximum similarity with only one difference. Also shown is the Pearson correlation coefficient (r) and its associated (one-tailed) p-value.

As can be seen in figure 4, the regression lines for the DPs and the control subjects are rather parallel for all conditions with houses suggesting that the effect of visual similarity did not impact differently on the DP group and the control group. In comparison, the slopes seem steeper for the DP group than for the control group in the conditions with faces, suggesting that increasing visual similarity impacted more on the performance of the DPs than the control subjects’ during processing of faces. To examine this more formally we first computed a difference-score between the performance of the DPs and the control subjects on similarity levels 0 and 3 respectively for each condition (1^st^ order, 2^nd^ order and the featural condition). We next examined whether the two difference-scores from each condition differed significantly from each other by means of Mann–Whitney *U* Exact tests. Hence, what is basically tested is whether there is an interaction between Group and Similarity level (focusing on the extreme ends of the similarity variable ignoring any difference at intermediate similarity levels 1 and 2). These analyses revealed no significant difference for neither the 1^st^ order condition (Z = −1.02, *p* = .39) or the featural condition (Z = −1.57, *p* = .12). However, in the 2^nd^ order condition, the difference between the DPs and controls was significantly larger for similarity level 3 than for similarity level 0 (Z = −2.27, *p* = .02). In summary, visual similarity did not seem to impact differently on the performance of the DPs and the controls with the exception of perceiving differences in 2^nd^ order relations for faces where the performance of the DP group was more affected than that of the control group as similarity increased.

#### 5.2.4. Single subject analysis

To examine the performance of the DPs individually, we compared the % correct responses on different-trials in each of the six conditions of the perceptual matching task of each DP with the mean % correct responses of the control participants using Crawford, Garthwaite and Porter’ method (2010). As can be seen in Table 5, where the results of these comparisons are summarized, four of the DPs (PP07, PP10, PP19 & PP27) scored within the normal range in in all conditions. In addition, one DP (PP04) performed within the normal range on all three conditions with houses. Four of the DPs (PP04, PP13, PP17 & PP18) exhibited significantly reduced performance with faces in one or more conditions, as did four DPs with performance with houses (PP09, PP13, PP17 & PP18). Hence, in terms of frequency, the DPs were just as impaired with houses as they were with faces.

As discussed in the introduction, good performance in terms of accuracy may be achieved at the price of prolonged RTs. With this in mind, we wanted to examine whether the four DPs, who performed within the normal range of the control group in all conditions, would also exhibit normal performance in terms of RT. Hence, we repeated the analyses presented above but now on the RTs of these DPs (PP07, PP10, PP19 & PP27) and their eight controls. We considered these analyses appropriate for this particular subsample of the DP group because they performed as accurately as the control subjects. We also included PP04 in these analyses when examining RT-performance with houses because PP04 had within normal-range performance in all three conditions with houses. Accordingly, 10 control participants were included in RT analyses with houses. As can be seen from table 6, only PP04 was significantly impaired in terms of RT. In conclusion, PP07, PP10, PP19 & PP27 performed within the normal range with both faces and houses with respect to both accuracy and RT.

**Table 5.**
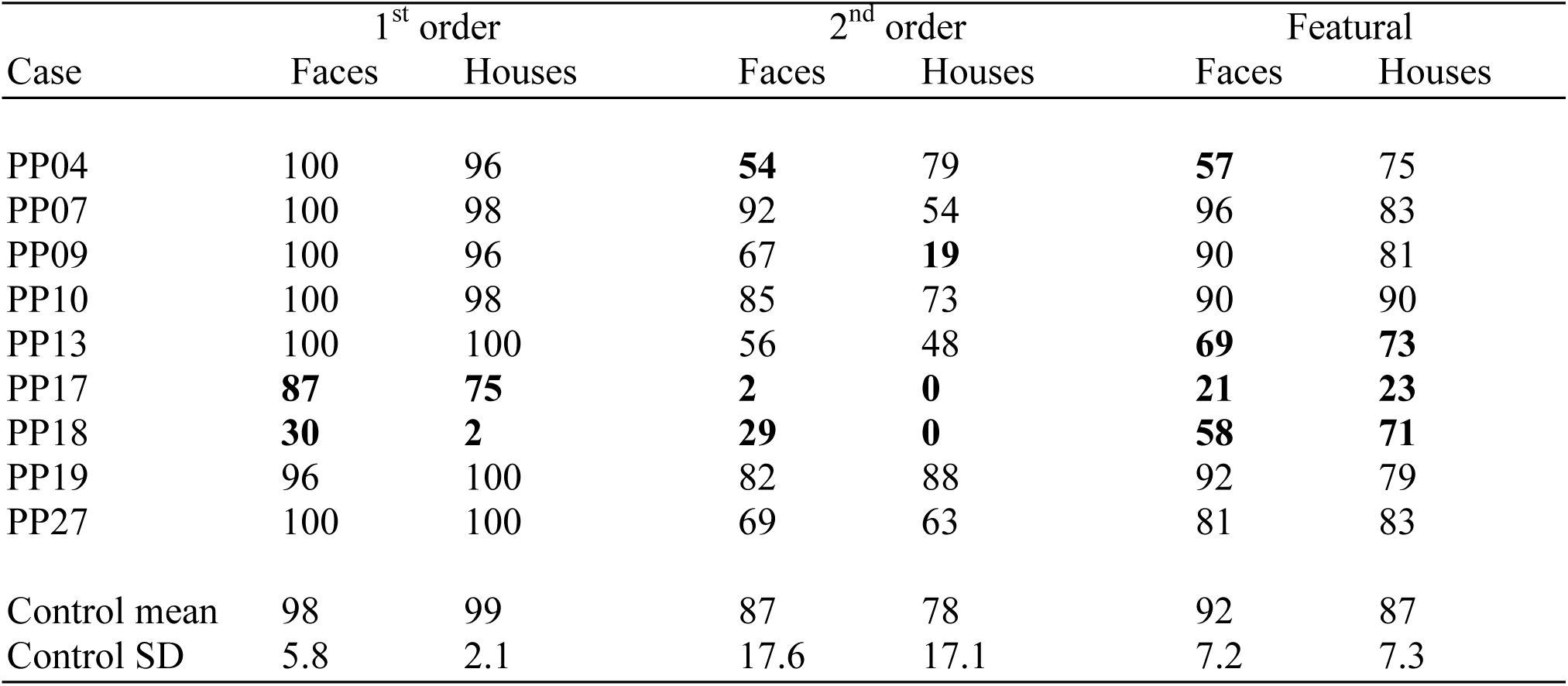
% correct responses to different-trials in the six conditions of the perceptual matching task for each of the individual DPs. Values in boldface designate performance deviating significantly from the mean of the matched control group as assessed by means of Crawford, Garthwaite and Porter’s (2010) method.

**Table 6.** Mean RT to different-trials in the six conditions of the perceptual matching task for the subsample of DPs with normal accuracy. Values in boldface designate performance deviating significantly from the mean of the matched control group as assessed by means of Crawford, Garthwaite & Porter’s (2010) method.

| Case | 1<sup>st</sup> order |  | 2<sup>nd</sup> order |  | Featural |  |
| --- | --- | --- | --- | --- | --- | --- |
|  | Faces | Houses | Faces | Houses | Faces | Houses |
| PP04 | ----- | 2426 | ----- | **4783** | ----- | **3677** |
| PP07 | 1447 | 1440 | 2826 | 3344 | 2515 | 2276 |
| PP10 | 1395 | 1607 | 2219 | 3066 | 2310 | 2454 |
| PP19 | 1533 | 2140 | 3520 | 2853 | 2669 | 2430 |
| PP27 | 1543 | 1596 | 1744 | 2779 | 1917 | 2142 |
| Control mean | 1294 | 1636 | 2161 | 2831 | 1961 | 2099 |
| Control SD | 277 | 426 | 692 | 913 | 644 | 605 |

### 5.3. Discussion

Considered as a group, the DPs generally exhibited poorer discriminability of both faces and houses compared with controls. This was reflected in their performance in all conditions except for 1^st^ order differences in faces where the DP group did not differ significantly from the control group in accuracy. In general, however, the DP group was not more affected by visual similarity than the control group in the 1^st^ order and featural conditions. Only when required to make discriminations based on differences in 2^nd^ order relations in faces was the DP group more affected by visual similarity than the control group. It is worth noting that this finding cannot reflect that this condition just happened to be the most challenging. While it was difficult, it was not more difficult than the 2^nd^ order condition with houses where the DP group and the control group were equally affected by increasing levels of visual similarity. Accordingly, at least this effect seems specific to faces.

Analyses of the individual performances of the DPs revealed a similar pattern as described for the group differences: Four DPs fell outside the range of the control subjects in one or more conditions with faces and five did so with houses. Except for one DP (PP09), who was only impaired with houses, it was the same DPs who fell outside the normal range with faces and with houses. Likewise, the four DPs who performed within the normal range in all conditions with faces also performed within the normal range with houses. Accordingly, there was no evidence of a face selective impairment in this task.

## 6. Testing for dissociations

Two of the four DPs (PP07 & PP27), who performed within the normal range with houses in the perceptual matching test also performed within the normal range on the object decision tasks and the Cambridge Car Memory Task. Hence, these subjects fulfil one of the premises for a dissociation. What further needs to be examined is whether the difference they exhibit in their performance with faces and objects is significantly greater than what can be expected in the normal population. To test this we adopted the approach suggested by Crawford et al. (2003) and Crawford and Garthwaite (2005) for testing of strong or classical dissociations in single-case studies (The revised standardized difference test; Crawford, Garthwaite & Porter (2010)). Specifically, we compared the difference in each DP’s score on the CFMT and their *A* sensitivity score in the object decision task with silhouettes; the condition in which the DPs as a group performed worse than the control group in terms of discriminability (*A*). These comparisons revealed that neither PP07 nor PP27 fulfilled the criteria for a classical or strong dissociation *(p* = .26 and *p* = .09 for PP07 and PP27 respectively). When the same analyses were conducted on the scores from the CCMT, similar results were obtained in that neither PP07 nor PP27 fulfilled the criteria for either a strong or classical dissociation *(p* = .06 and *p* = .37 respectively).

## 7. General discussion

If face recognition and object recognition rest on separate domain-specific operations, we would expect to find subjects with face recognition deficits who perform within the normal range with recognition of other categories of objects, and the reverse; a double-dissociation. In the present study we examined one side of such a potential double-dissociation. This was done by testing whether a relatively large group of subjects (*N* = 10) with developmental prosopagnosia (DP) performed within the normal range on demanding visual object processing tasks. While we do find that our group of DPs performed within the normal range on a demanding object recognition task with regular drawings, we also find that they are impaired relative to control participants with degraded stimuli (silhouettes and fragmented forms) and with within-class recognition of objects (cars). In addition, the DPs’ object recognition performance with degraded stimuli is systematically related to the severity of their face recognition deficit. Hence, we find that their face recognition performance is highly correlated with recognition of both silhouette objects (*r* = .87) and fragmented objects (*r* = .78).

So, is there no evidence of face selectivity in our sample of DPs? If one considers the DPs individually there appears to be some evidence because many of them performed within the normal range on many of the tasks. Indeed, two of them (PP07 & PP27) scored within the normal range on *all* tasks including the ones with degraded stimuli, which is quite impressive. However, when judged on a group level it is also the case that the DPs were in fact significantly impaired in the majority of the tasks performed: Within-class recognition of objects (cars), object decision with degraded material, and perceptual matching of faces and houses (2^nd^ order and featural differences). Hence, we are caught in an old dilemma: Should (neuropsychological) evidence be based on individual cases, which may leave something to be desired in terms of representability and statistical power, or should it rather be based on group studies which typically excel in both respects but which may yield results that are basically “averaging” artefacts (Caramazza, 1986)? And how do we approach the old dogma that dissociations trumps associations, in cognitive neuropsychology, when we can show significant (and impressive) correlations on a group level, while some individuals exhibit borderline significant dissociations? The most reasonable thing to do is to weigh the evidence, considering the strength and weaknesses of each approach.

In doing so we first note that even though not all DPs performed significantly outside the normal range when considered individually, especially not with fragmented forms and within-class recognition of objects, most of them did perform in the lower normal range in terms of either accuracy (discriminability) or RT. Was this not the case, the reliable group differences would not have been found. Secondly, even the two DPs who performed the best (PP07 & PP27) did not fulfil the criteria for a classical or even strong dissociation. Although PP27 did come close to showing a dissociation when considering the difference in his performance on the CFMT and the object decision task with silhouettes *(p* = .09), the difference in his performance between the CFMT and the CCMT was not significantly different from what can be expected in the normal population *(p* = .37). The opposite was true of PP07, the other of the best performing DPs, where the difference between the CFMT and CCMT approached significance *(p* = .06), whereas the difference between the CFMT and the object decision task with silhouettes did not *(p* = .26). Thirdly, there was quite a lot of variability in our control sample, possibly owing to differences in e.g., age and educational level. While this is not a problem when comparing DPs and control participants as groups, precisely because they were matched on these parameters, it becomes more problematic when the individual DPs are compared with the (mean and SD of the whole) control sample, as is done in the single subject analyses. Finally, the systematic relationships between performance with degraded material and face recognition performance strongly suggests that the results of the group comparisons do not reflect averaging artefacts. In our opinion this finding is important because a correlation, as opposed to a dissociation, does not depend on the performance of a control group; a performance which may vary from sample to sample (cf. section 7.1 below). With these considerations in mind, we do not think that our findings yield support for face selective impairments in our sample of DPs. On the contrary, our data suggests that their object recognition performance is also impaired, although often in a subtle manner. Indeed, the finding that the severity of their object recognition deficit is systematically related to the severity of their face recognition deficit is difficult to account for should their face deficit be selective.

If our findings are representative of DP in general, our conclusion regarding non-selectivity contrasts with the conclusion reached in several other studies reporting a dissociation between face and object recognition ability. To our knowledge, the most compelling evidence for a dissociation involving impaired face recognition and normal object recognition has been reported by Duchaine and Nakayama (2005). They compared the performance of seven developmental prosopagnosics (DPs) on old/new recognition memory tests involving the following categories: Faces, Cars, Houses, Scenes, Horses, Tools, and Guns. Of these DPs, two subjects (cases M1 and M2) performed within the normal range on all contrast categories when performance was assessed by means of discrimination sensitivity, and one subject (case F2) performed within the lower normal range on all categories. However, none of these DPs performed within the normal range in terms of RT. Cases M1 and M2 – who were the best performing of the DPs in terms of accuracy on the contrast tasks – fell more than two SDs below the mean on respectively all or four of the contrast categories, whereas case F2 fell more than two SDs below the mean on two of the contrast categories. Hence, if both accuracy and RT are considered, none of the DPs reported by Duchaine and Nakayama (2005) performed within the normal range with objects. The same is true of other studies where the performance of DP-subjects fall outside the normal range on either RT or accuracy (e.g., Garrido, Duchaine and Nakayama, 2008; Duchaine, Yovel, Butterworth, & Nakayama, 2006).

Even though we advocate for an association between face and object recognition impairments in DP, which might lead to the conclusion that faces are not ‘special’, there is no doubt in our minds that the DPs included in our study experience tremendously greater problems with faces than they do with objects in everyday life. In this sense faces are special. To make our stance clear, we find no compelling evidence in the literature on DP or in the present study suggesting that face recognition is so special that it can be selectively impaired, but it is special enough to place greater strains on the recognition system – or parts of the system – than recognition of other categories of objects typically do, causing recognition of faces to suffer significantly more. Accordingly, it is only when the recognition system is seriously challenged that recognition deficits with other objects than faces becomes apparent. We also note that our claim regarding a lack of convincing cases of selectivity (dissociations) in face recognition only applies to the field of DP. While the same may be true for acquired prosopagnosia (for a discussion of this see Busigny, Joubert, Felician, Ceccaldi, & Rossion, 2010), a detailed discussion of this evidence falls outside the scope of the present paper.

### 7.1. A concluding note on selectivity

But what is “selectivity” anyway? In a sense, selectivity is a result of how observations are treated. The traditional way, which we also adopted here, is to classify an individual’s performance as abnormal if it falls outside the range of “normal” participants; typically defined as 2 SDs below the mean of the normal subjects. If it does not, the individual’s performance is considered normal. To some extent this cut-off is arbitrary and its application will yield different outcomes depending on the norms it is applied to. However, if we abandon the use of cut-off scores, selectivity ceases to exist. Then it becomes a question of degree rather than kind. We are not arguing that a cut-off at 2 SDs necessarily represents an insensible way to operationalize a deficit. Our point is rather that it is ‘just’ an operationalization which does not (necessarily) imply that the cognitive or neural machinery which supports recognition must consist of two separate systems (modules) if it can be demonstrated that some subjects show “> 2SDs selectively” impaired face recognition, whereas others show “> 2SDs selectively” impaired object recognition.

If we look at things from this perceptive it seems productive to further investigate, for example, what it is that recognition of degraded objects has in common with face recognition. One possibility is that it is related to the derivation of global shape information, which has been suggested to play a pivotal role in recognition of both silhouettes and fragmented forms (Gerlach & Toft, 2011), as well as face recognition (Sergent, 1986).

## Acknowledgments

We wish to pay our gratitude to the Friends of Fakutsi Initiative (FFI).

